# Studying metal-protein interactions using fluorescent protein indicators

**DOI:** 10.1101/2022.10.13.512174

**Authors:** Joel C. Rosenbaum, Anne E. Carlson

## Abstract

Metals are widespread environmental toxins that disrupt normal cellular processes through their interactions with proteins and other macromolecules. In this study, we developed the metal-sensitive fluorescent protein mseGFP as a ratiometric reporter capable of binding heavy metals. We found that mseGFP bound mercury and lead tightly but had substantially lower sensitivity to other metals. By comparison, the redox sensor roGFP2 functioned as a ratiometric indicator for transition metals, with the highest sensitivity for copper, followed by nickel and cobalt. mseGFP and roGFP2 could also report metal binding through fluorescence quenching, and we used this effect to measure high affinity interactions for both proteins with copper and iron. Crystal structure analysis of mseGFP complexed with phenylarsine oxide revealed an unexpected mode of heavy metal interaction, with mseGFP binding PAO with 2:2 stoichiometry. Glutathione strongly inhibited most metal interactions with the fluorescent protein reporters, but increased the affinity of arsenic and cadmium for mseGFP. When expressed in HEK293T cells, mseGFP reported uptake of mercury and phenylarsine oxide from surrounding media. Glutathione depletion enhanced binding of phenylarsine oxide to mseGFP in cells, validating the importance of glutathione in modulating metal-protein interactions.

## Introduction

Metals are widespread and persistent environmental toxins with profound impacts on public health. Given their abundance and potency, heavy metals such as arsenic, lead, mercury, and cadmium are considered among the most hazardous toxins in the lived environment(1,2). Toxic exposure to heavy metals affects more than one hundred million people globally(3) and is linked to a broad spectrum of illnesses, including cardiovascular disease(4), cognitive disorders(5), and cancer(6,7). Even less toxic metals, including essential micronutrients such as copper, iron, and zinc can also produce adverse effects from excessive exposure or diseases that disrupt homeostasis(8). Having reporters for these metals is vital for understanding how metals interact with cellular factors and for studying processes that affect metal uptake and homeostasis(9).

Toxic metal exposure harms cells through various mechanisms, including by producing reactive oxygen species and disrupting enzyme function(10). For example, some metals can generate free radicals through catalysis of peroxide(11,12) or by activating oxidizing enzymes(13). Metals can also exhaust cellular antioxidants by inhibiting redox enzymes such as thiol oxidoreductases(14,15). Generally, metals can disrupt normal enzyme functions involved in a wide range of physiological processes, including heme biosynthesis, oxidative phosphorylation, and DNA repair(2). Heavy metals are particularly disruptive to enzymes with cysteine-containing motifs as heavy metals have a strong affinity for thiols(16).

Metals can also interact with abundant cellular small molecules that have metal coordination properties. These low molecular weight ligands alter metal interactions with proteins and other cellular factors by catalyzing ligand exchange(17,18) or through the formation of ternary complexes that alter their mode of binding(19). Glutathione is one of the most abundant low molecular weight ligands, with millimolar cellular concentrations(20). Reduced glutathione (GSH) can form complexes with transition and heavy metals, consequently altering their interactions inside the cell. Glutathione is protective against metal stress(21,22), and this effect is generally attributed to its role as a cellular antioxidant. However, glutathione complexation may also provide direct protection from heavy metals by facilitating export(23). Despite the abundance of glutathione in cells, its effect on metal-protein interactions has only been examined in select cases(17).

We used fluorescent proteins to measure metal interactions with proteins featuring surface proximal cysteines. These include the reduction-oxidation-sensitive GFP (roGFP2) previously developed to measure cellular redox levels(24,25) and a related metal-sensitive GFP (mseGFP) explicitly designed for this study. We report metal interactions with both proteins *in vitro* and with mseGFP in mammalian cells. Finally, we determined the influence of GSH on these interactions, demonstrating that physiological concentrations of glutathione dramatically alter metal interactions with proteins.

## Results

We created a fluorescent metal reporter, <u>m</u>etal-<u>s</u>ensitive GFP (mseGFP), based on the design of the redox-sensitive fluorescent protein roGFP2(24,25). roGFP2 is a redox reporter protein with proximal cysteines (C147, C204) placed with a Cα distance of 4.4-4.8Å. Under oxidizing conditions, the formation of a strained disulfide bond causes a conformational change that alters the fluorescence properties of roGFP2(25). We used a cysteine pair (C147, C202) in mseGFP with a Cα distance of 7.3Å, far enough apart to preclude the formation of disulfide bonds, but still close enough to allow for a similar conformational change resulting from metal binding (Fig. 1A).

**Figure 1.**
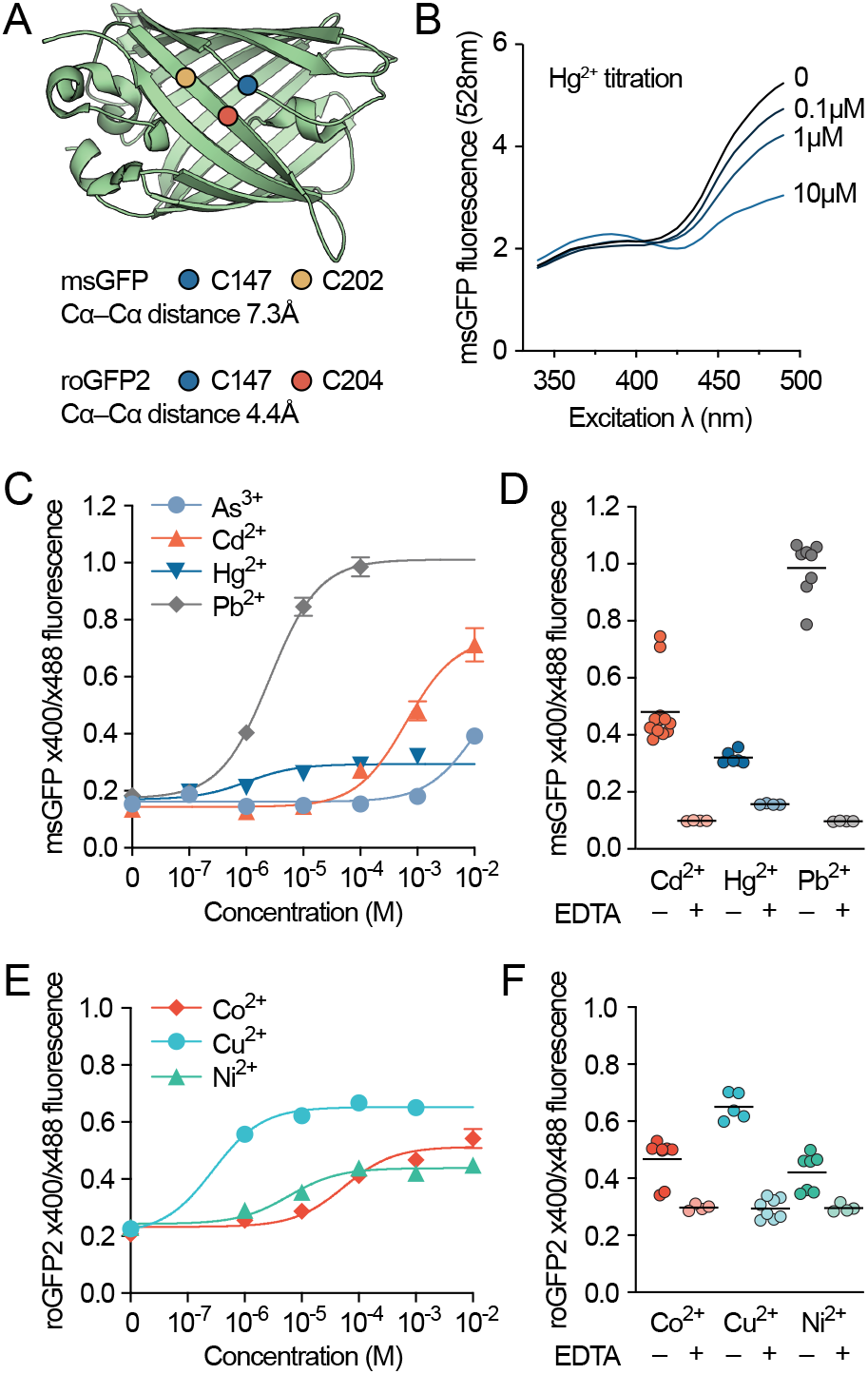
Measuring metal-protein interactions with fluorescent reporters. *A*, engineered cysteine sites for mseGFP and roGFP2 mapped on reference GFP structure (PDB 4EUL). *B*, excitation spectra of mseGFP during titration of Hg^2+^. Left axis indicates relative fluorescence of emission measured at 528nm. *C*, ratiometric change in mseGFP fluorescence during titrations of indicated metals. Error bars indicate SEM. *D*, reversal of metal-induced (1 mM for Cd^2+^ and 100μM for Pb^2+^ and Hg^2+^) ratiometric fluorescence changes in mseGFP by addition of 5 mM EDTA. *E*, same as (*C*), but for roGFP2. *F*, reversal of metal-induced (1 mM for all metals shown) fluorescence changes in roGFP2 by addition of 5 mM EDTA.

We performed titrations with mseGFP and metals with known thiol-binding characteristics. We first determined that Hg^2+^ induced a ratiometric change in the excitation spectra of mseGFP (Fig. 1B) similar to the effect produced by oxidation on roGFP2(24). All tested heavy metals (Pb^2+^, Hg^2+^ Cd^2+^, As^3+^) induced observable changes in ratiometric mseGFP fluorescence, with Pb^2+^ (Kd = 2.6 μM) and Hg^2+^ (Kd = 1.0 μM) having the highest potency. Cd^2+^ (Kd = 640 μM) and As^3+^ (Kd > 10 mM) had a more modest affinity as measured by change in mseGFP fluorescence (Fig. 1C). For each of these metals except As^3+^, fluorescence changes were reversed upon addition of 5 mM EDTA (Fig. 1D). By comparison, transition metals (Fe^2+^, Fe^3+^, Ni^2+^, Co^2+^, Cu^2+^) did not produce a ratiometric change in mseGFP fluorescence. The post-transition metal Zn^2+^ produced a change in fluorescence comparable to that induced by As^3+^ (Fig. S1A).

In titrations with metals and roGFP2, the transition metals Cu^2+^ (Kd = 0.3 μM), Ni^2+^ (Kd = 5.9 μM), and Co^2+^ (Kd = 55.4 μM) produced changes in ratiometric fluorescence (Fig. 1E), whereas heavy metals did not (Fig. S1B). The transition metal-induced changes in ratiometric roGFP2 fluorescence were reversible by EDTA and not β-mercaptoethanol (Fig. 1F, S2), indicating that this effect is caused directly by metal binding and not by disulfide formation(26,27). Addition of H_2_O_2_ induced a ratiometric change in roGFP2 fluorescence at high concentrations, consistent with previous results(24), but had no effect on mseGFP fluorescence (Fig. S1C).

We observed fluorescence quenching for both mseGFP and roGFP as metals were added. This effect was particularly noticeable with transition metals, which form complexes that absorb light in the visible spectrum. We hypothesized that this was due to quenching by FRET with the GFP chromophore, previously observed in fluorescent proteins with engineered metal binding sites(28,29). FRET is a distance-dependent, non-radiative form of energy transfer whose efficiency also depends on the overlap between the emission of the donor fluorophore and the absorbance of the acceptor chromophore(30). We observed the strongest quenching with transition metals (Cu^2+^, Fe^2+^, and Fe^3+^) whose absorbance spectra overlapped with the identical emission spectra of roGFP2 and mseGFP (Fig. 2A, left). FRET radius (R_0_) values calculated for these fluorescent proteins with Cu^2+^ (13.3Å), Fe^3+^ (15.1Å), and Fe^2+^ (9.1Å) support the possibility of FRET-dependent quenching, as these distances exceed the GFP chromophore-C147 side chain distance of 7.5Å in roGFP2(25) (Fig. 2A, right). Quenching for mseGFP (excited at 488 nm) clearly followed the expected pattern for single site binding (Fig. 2B), with Cu^2+^ (Kd = 0.5 μM) and Fe^3+^ (Kd = 1.2 μM) binding more tightly than Fe^2+^ (Kd = 120 μM). For Cu^2+^ and Fe^3+^, quenching was abolished by the inclusion of 5 mM EDTA in the assay buffer, signifying that quenching was the result of direct interaction between these metals and mseGFP. Some quenching was still observed for Fe^2+^ even in the presence of excess EDTA, with this possibly caused by the inner filter effect(31). In titrations with roGFP2, we observed similar fluorescence quenching curves for Fe^3+^ (Kd = 2.2 μM) and Fe^2+^ (Kd = 120 μM) as with mseGFP (Fig. 2C). However, Cu^2+^ appeared to quench roGFP2 with lower affinity (Kd = 8.0 μM) compared to mseGFP, although the binding model does not account for ratiometric changes in roGFP2 fluorescence (Fig. 1D).

**Figure 2.**
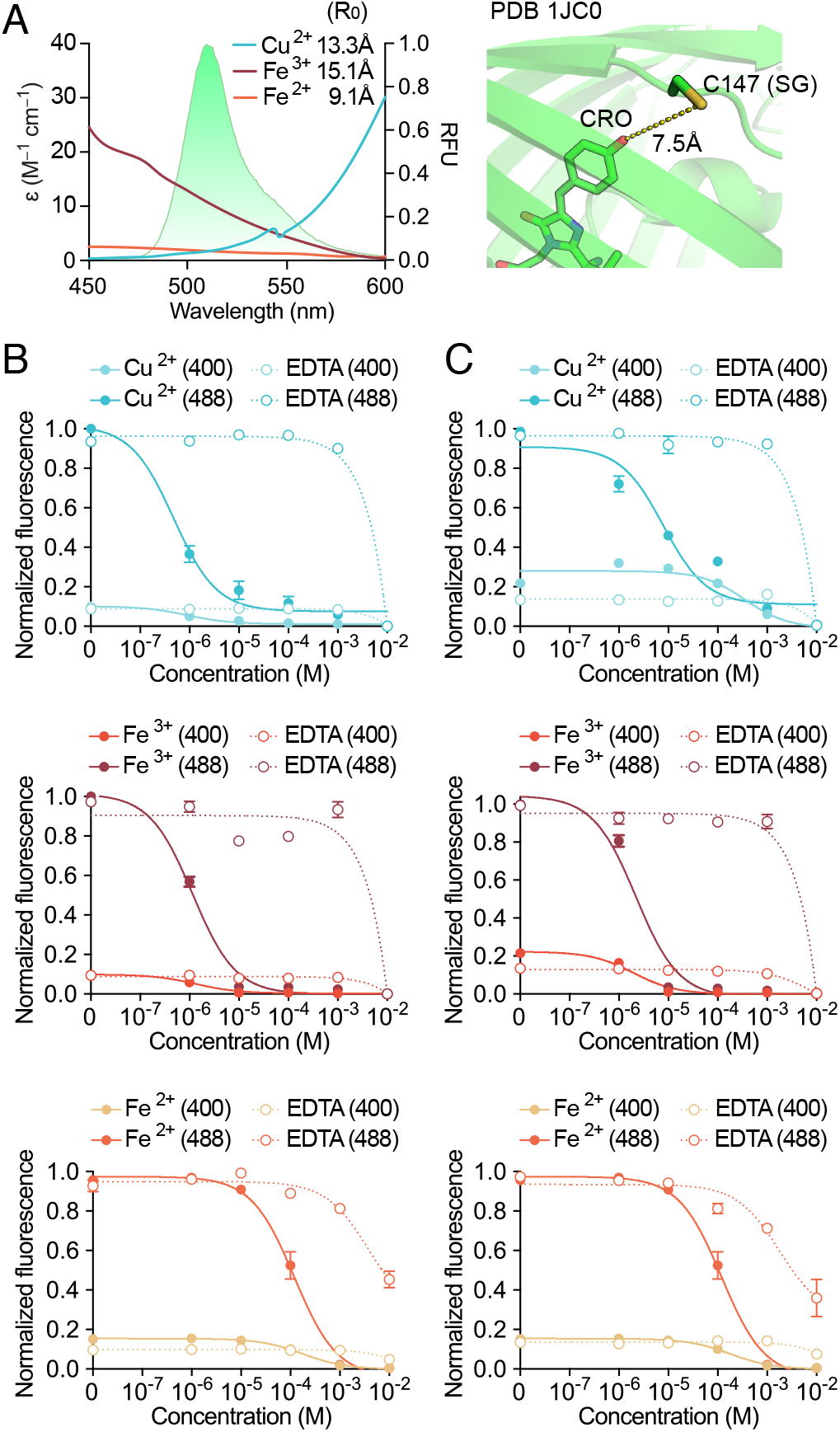
mseGFP and roGFP2 quenching by metal chromophores. *A*, overlay of FP emission spectra (identical between mseGFP and roGFP2) with absorption spectra of coordinated Cu^2+^, Fe^3+^, and Fe^2+^ complexes. Also shown (inset) is the calculated Forster radii (R_0_), and (right) the distance between the phenolate of the GFP chromophore and the thiol of C147 in roGFP2 (PDB 1JC0). *B*, quenching of mseGFP fluorescence with indicated metals in (*A*) along with its reversibility by inclusion of 5mM EDTA. Error bars indicate SEM. *C*, same as (*B*) but for roGFP2.

We observed that As^3+^ bond to mseGFP with weak affinity (Fig. 1A), although it had been previously reported to bind similar sites more tightly(32). We sought to determine how arsenic speciation might affect this interaction. Inorganic arsenic has two valence states, with trivalent As^3+^ interacting with thiol groups in proteins(33). Beyond that, a diverse assortment of organoarsenical compounds are also found in the environment, having been formed through biological or chemical synthesis(34). Consistent with previous studies, the pentavalent As^5+^ did not bind mseGFP (Fig. 3). However, the organoarsenical compound phenylarsine oxide (PAO) bound mseGFP with an affinity (Kd = 2.4 μM) comparable to that of Pb^2+^ and Hg^2+^ (Fig. 1B) and exceeding the affinity for As^3+^ by nearly four orders of magnitude. These data suggest that substitution might increase the affinity of As^3+^ for protein thiols (Fig. 3B).

**Figure 3.**
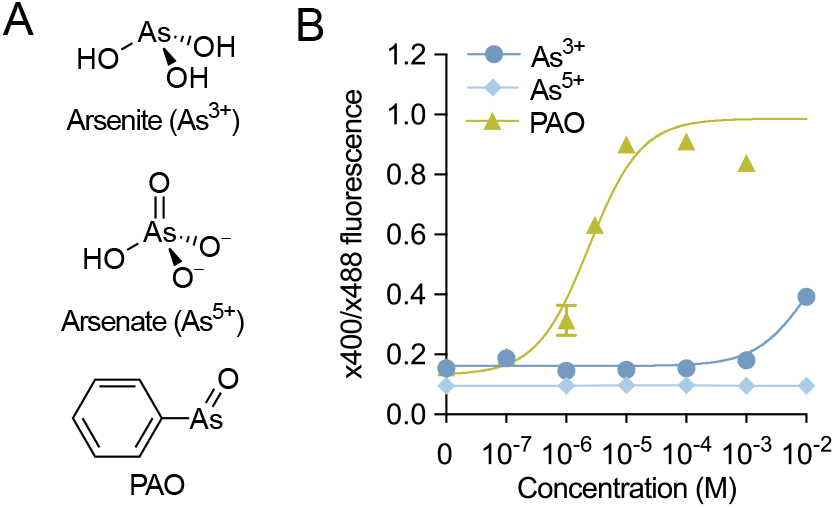
Arsenic speciation affects reactivity with mseGFP. *A*, ionic form of arsenic species used in this assay (pH 7). As^3+^ data reproduced from Fig 1. *B*, ratiometric change in mseGFP fluorescence during titrations of indicated arsenic species. Error bars indicate SEM.

We next evaluated the effect of glutathione on metal-mseGFP interactions. Glutathione is the central redox agent and most abundant thiol in most aerobic organisms(35). In mammalian cells, glutathione concentrations range from 1-10 mM(20), with the reduced form comprising ~98% of the cellular pool(36). Including 1 mM reduced glutathione (GSH) included in the assay dramatically altered the metal-mseGFP affinity of every tested metal (Fig. 4). We observed a significant increase in the affinity of As^3+^ for mseGFP (Fig. 4A, left) likely resulting from the formation of labile As(GS)3 complexes that readily react with proteins(37,38) (Fig. 4A, right). Overall, we observed significant increases in mseGFP binding affinity for As^3+^ and Cd^2+^ and a modest increase for Zn^2+^. By comparison, we saw decreases in affinity for all other metals tested (Fig. 4B). When included in titrations with roGFP2, 1 mM glutathione decreased the affinity of all metals except for Co^2+^, which was not affected (Fig. 4C). For both proteins, the effect of glutathione on metal binding was usually substantial, often changing affinities by an order of magnitude or more (Fig. 4B, C). We also observed that the effect of glutathione was concentration-dependent, with various concentrations within the range of physiological variation having distinct effects on metal binding. For As^3+^ and GSH into mseGFP revealed a GSH-dependent increase in As^3+^ binding at GSH concentrations up to 3 mM, followed by a decrease at 10 mM GSH (Fig. 4D). By comparison, any addition of GSH reduced the affinity of PAO for mseGFP (Fig. 4D). Titrations for Cd^2+^ and Zn^2+^ with GSH also revealed GSH-dependent affinity increased for mseGFP, although Cd^2+^ precipitated with higher concentrations of GSH(39) (Fig. 4D).

**Figure 4.**
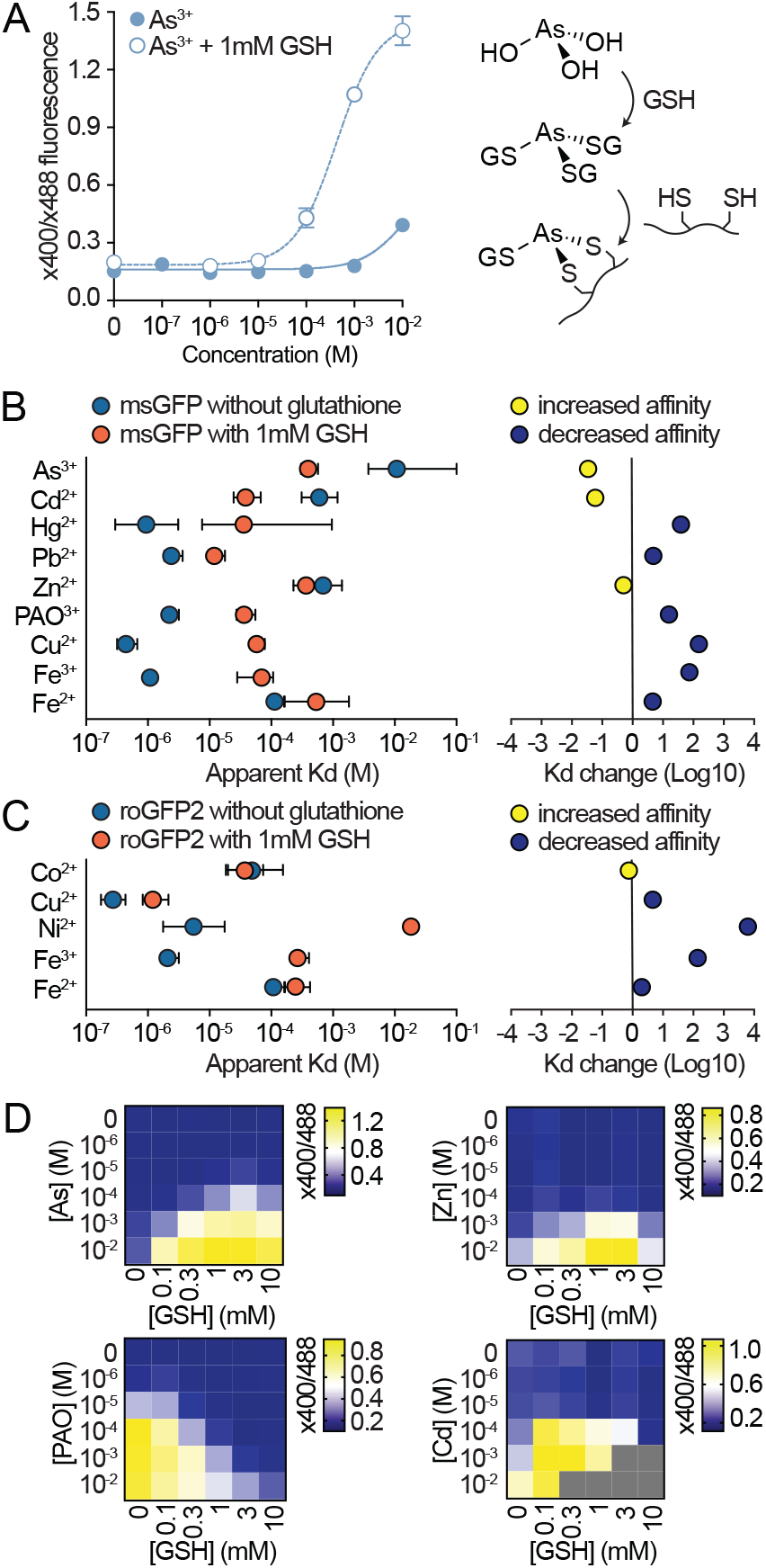
Glutathione alters interactions between metals and fluorescent reporters. *A*, ratiometric fluorescence of mseGFP in a titration of As^3+^ with or without 1 mM glutathione. SEM indicated by error bars. Hypothesized mechanism (right) of glutathione mediated As^3+^ interaction with cysteine pairs. *B*, Summarized effects of 1mM glutathione on the interactions of mseGFP with indicated metals. Error bars indicate the confidence intervals of Kd estimates. Affinity changes (right) between mseGFP and indicated metals after addition of glutathione. *C*, Same as for (*B*), but with roGFP2. D, mseGFP titrations with As^3+^, Zn^2+^, PAO, and Cd^2+^ against glutathione. Scale for ratiometric titration is shown (right) for each array. Grey boxes indicate samples that could not be acquired due to precipitation.

Next, we determined the efficacy of mseGFP in reporting metal binding within cells. Surprisingly, exposing HEK293T cells stably expressing mseGFP to micromolar concentrations of metals largely had no effect on mseGFP fluorescence (Fig. 5A-B). However, Hg^2+^ and PAO induced ratiometric changes in cellular mseGFP fluorescence consistent with results obtained *in vitro* (Fig. 5B, 3B). We found that PAO-induced changes in mseGFP fluorescence occurred more rapidly than observed for Hg^2+^, likely due its increased cell permeability. However, we found that the effect of Hg^2+^ was stronger (Fig. 5B). We then determined the effect of glutathione depletion on mseGFP binding with PAO, interactions for which glutathione had large effects *in vitro* (Fig. 4B). Pretreatment of mseGFP-expressing HEK293T cells with the glutamate-cysteine ligase inhibitor(40) buthionine sulfoximine (BSO, 100 μM), increased the effect of PAO on mseGFP fluorescence (Fig. 5C-D).

**Figure 5.**
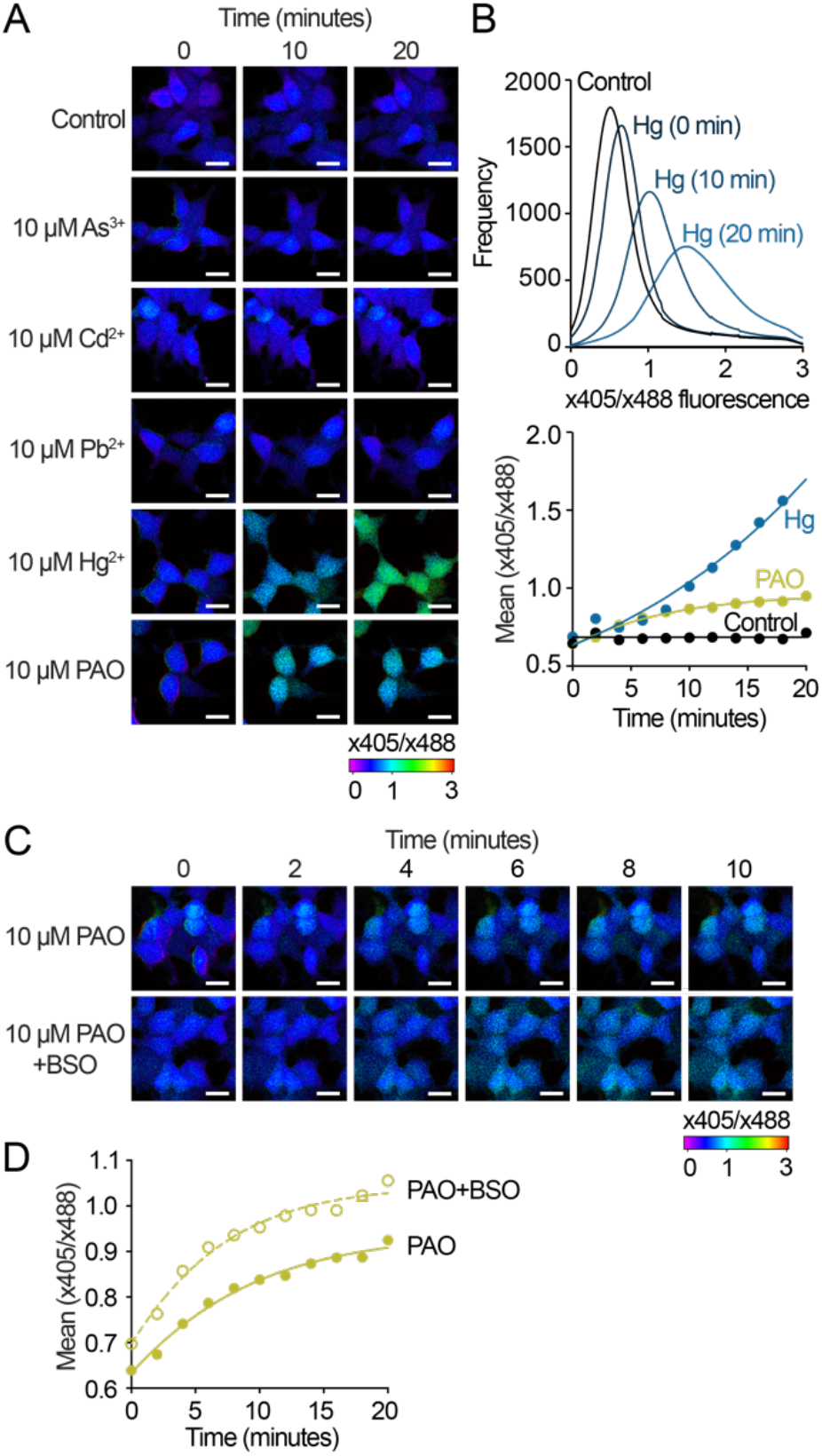
Heavy metals alter mseGFP fluorescence in HEK293T cells. *A*, ratiometric fluorescence of mseGFP in HEK293T cells after exposure to heavy metals. Expanded timecourses (at right) shown for metals that produced changes in mseGFP fluorescence. 20 μm scale bars are shown for reference. *B*, the ratio of violet-(x405) to blue-(x488) excited fluorescence changes over time for mseGFP-expressing cells exposed to Hg^2+^ (top). Mean x405/x488 ratio plotted over time for each experimental condition (bottom). *C*, glutathione depletion with BSO (100 μM) increased the rate of PAO-induced effects on mseGFP fluorescence in HEK293T cells. 20 μm scale bars are shown for reference. *D*, mean x405/x488 ratio charted for BSO- and untreated cells after addition of PAO.

We also attempted structural determination of mseGFP-metal complexes with crystallization screens of mseGFP and interacting metals. Among these trials, mseGFP-PAO produced crystals that diffracted to 1.89Å. Following molecular replacement and refinement, we produced a crystallographic model that surprisingly placed PAO at the interface between subunits of an antiparallel mseGFP dimer in a 2:2 conformation, linking C147 of one subunit to C202 of the other (Fig. 6A). Our composite OMIT (2mFo-DFc) map(41) supported the placement of PAO at this position (Fig. 6B). In contrast to previous studies comparing the structural arrangement of oxidized (1JC1) and reduced (1JC0) forms of roGFP2(25), we were unable to identify any significant structural changes in PAO-bound mseGFP compared to reference eGFP(42) (Fig. 6C).

**Figure 6.**
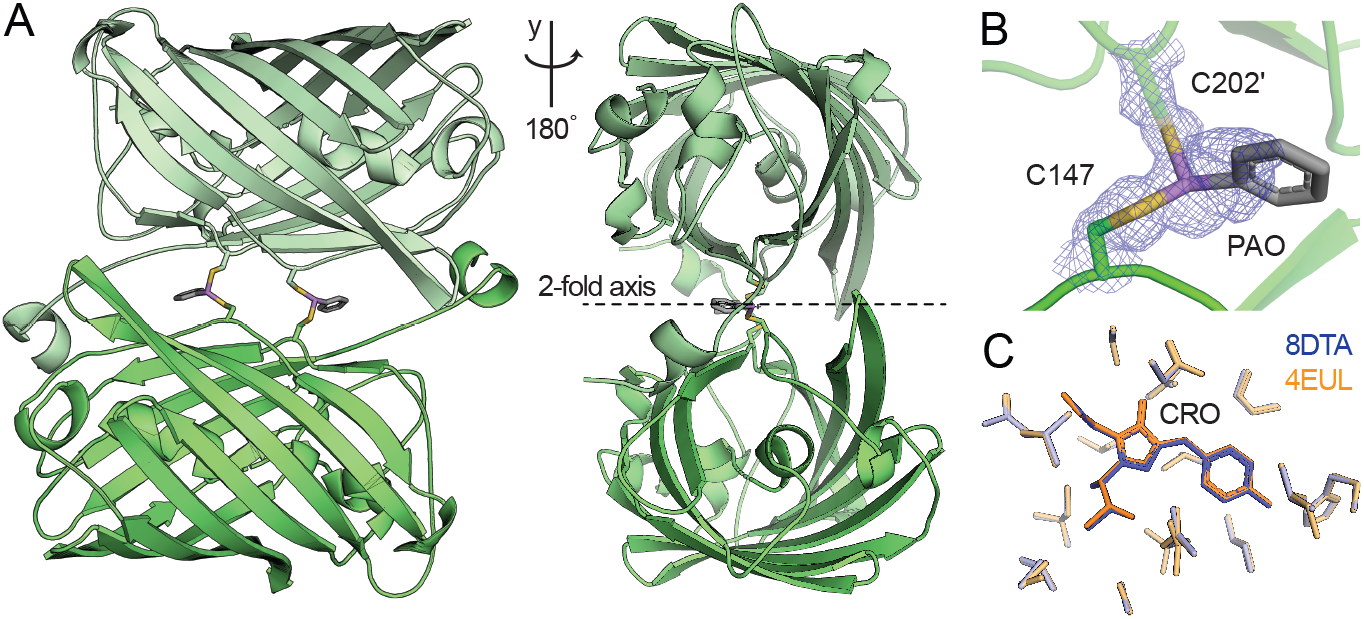
Crystallographic structural model of mseGFP-PAO complexes. *A*, model of mseGFP-PAO complexes resolved by x-ray crystallography. An mseGFP monomer (dark green) complexes with PAO in a 2:2 stoichiometry with its symmetry mate (light green). Rotated view (right) of the mseGFP-PAO complex showing the 2-fold rotation axis. *B*, composite OMIT map of one PAO coordinated by C147 and C202 across two mseGFP monomers. *C*, superposition of mseGFP-PAO (PDB 8DTA) on reference GFP (PDB 4EUL) showing that the chromophore environment is largely unchanged between the two structural models.

## Discussion

In this study, we engineered the fluorescent protein mseGFP from roGFP2, and compared the ability of both fluorophores to report interactions between metals and proteins. We noted that roGFP2 exhibited a previously unreported sensitivity to transition metals, with a reversible change in ratiometric fluorescence caused by titrations of Ni^2+^, Co^2+^, and Cu^2+^ that fit a single site binding model (Fig. 1B). Cu^2+^, but not Ni^2+^ or Co^2+^, also bound mseGFP as determined by fluorescence quenching (Fig. 2B). Interestingly, heavy metals induced a measurable ratiometric change in mseGFP, but had no observable effect on roGFP2 (Fig. 1, S1). Since mseGFP differs from roGFP2 only in the distance between its metal-binding cysteine residues (Fig. 1A), these variations in metal ion selectivity may be explained by differences in bond lengths and coordination geometries(43,44). This potentially also explains the discrete Bmax values observed for each metal (Fig. 1B, 1C).

We observed fluorescence quenching in both reporters for several of the metals tested. The strongest quenching was observed for the colored transition metals Cu^2+^, Fe^2+^, and Fe^3+^, whose absorbance overlaps with the identical emission spectra of mseGFP and roGFP2 (Fig. 2A). The spectral properties of Cu^2+^ have been exploited in measuring short-range interactions by FRET(45–47). FRET with an iron acceptor has similarly been observed in experiments linking a heme protein with eGFP(48). One surprising finding in our studies was that Cu^2+^ had a greater quenching effect on mseGFP compared to roGFP2 (calculated transfer efficiency of 0.94 for mseGFP compared to 0.80 for roGFP2) (Fig. 2), even though one of the coordinating cysteines (C202 for mseGFP, C204 for roGFP2) is located further from the GFP chromophore. Transition metals lack the orientation dependence of organic chromophores for FRET(47) and quenching efficiency should depend entirely on the distance between the donor fluorophore and the acceptor Cu^2+^ ion. The fact that mseGFP and roGFP2 do not follow this pattern raises the possibility that other quenching processes may be involved in Cu^2+^-mediated quenching of mseGFP.

Cells contain pools of small molecules that modulate metal interactions with proteins(17). Among the most abundant of these is glutathione, which in its reduced form is the predominant thiol in aerobic organisms(20,35). We found that GSH inhibited most interactions between metals with either mseGFP or roGFP2 (Fig. 4). However, GSH increased the affinity of As^3+^ and Cd^2+^ for mseGFP, likely through the formation of ternary complexes(39). In the presence of excess glutathione, arsenite forms As(GS)3 complexes that subsequently bind proteins through thiol exchange(37,38). We found that GSH enhances As^3+^ binding to mseGFP in a concentration-dependent fashion up to 3 mM, with higher concentrations of GSH exerting an inhibitory effect (Fig. 4D). By contrast, GSH at any concentration inhibited the interaction between the organoarsenical PAO and mseGFP. These findings suggest that, as with other metals, GSH also acts as a competitive inhibitor of arsenic-protein interactions. This is also likely true for Cd^2+^, which showed a similar GSH-dependent increase in affinity as concentrations exceeded 100μM (Fig. 4D). We found that binding of Zn^2+^ to mseGFP was enhanced by addition of GSH, although this effect was more modest than those observed for As^3+^ and Cd^2+^. Our findings are in line with previous observations that GSH alters the kinetics of Zn^2+^ binding to fluorescent indicators(19,49).

A surprising finding in our study is how mseGFP bound metals when expressed in HEK293T cells. Some metals that strongly bound mseGFP *in vitro*, such as Pb^2+^, had almost no effect on cellular mseGFP fluorescence (Fig. 6A). By comparison, Hg^2+^ and PAO had a rapid and profound effect on mseGFP fluorescence (Fig. 6A). We found that depleting glutathione by pretreatment with BSO had effects in agreement with our *in vitro* assays with PAO (Fig. 4D). As expected, pretreatment with BSO increased PAO-induced changes in mseGFP fluorescence (Fig. 6B). Considered with the results of our *in vitro* experiments, these results indicate that glutathione is an important factor alongside transporters, metallothioneins, and other low molecular weight ligands, that plays a determinative role in metal-protein interactions in cells.

## Experimental procedures

### Reagents

Copper (II) chloride dihydrate (Cu^2+^), cobalt (II) chloride hexahydrate (Co^2+^), iron (II) sulfate heptahydrate (Fe^2+^), iron (III) chloride (Fe^3+^), nickel (II) chloride hexahydrate (Ni^2+^), zinc (II) sulfate heptahydrate (Zn^2+^), cadmium (II) chloride hemipentahydrate (Cd^2+^), lead (II) nitrate (Pb^2+^), mercury (II) chloride (Hg^2+^), and hydrogen peroxide (H2O2, 30% in water) were purchased from Fisher Scientific. Buthionine sulfoximine (BSO), sodium arsenate (V) (AsO_4_^3-^, herein described as As^5+^), sodium arsenite (III) (AsO_3_^3-^, herein described as As^3+^), and phenylarsine oxide (PAO) were purchased from Millipore Sigma. Except for phenylarsine oxide, metal stocks were prepared at 100 mM in water, with the pH adjusted to 7 with 1M NaOH. Phenylarsine oxide stocks were prepared in DMSO.

### Plasmid constructs

pQE-60 roGFP2-His was a gift from Tobias Dick (Addgene #65046)(50). The construct for mseGFP was generated by substitution of codons encoding two amino acids (S202C, C204C) from parent roGFP2. Both mseGFP and roGFP2 were cloned into the bacterial expression plasmid pKM260. mseGFP was also cloned into pcDNA3 (Invitrogen) for mammalian expression. All cloning was performed by NEB HiFi assembly (New England Biolabs) and validated by whole plasmid sequencing (Plasmidsaurus).

### Protein expression and purification

pKM260-mseGFP and pKM260-roGFP2 were transformed into BL21(DE3)-pRARE2 cells by electroporation (0.1 mM gap, 2,000 V) at 4°C and plated on LB plates containing 100 μg/mL of ampicillin. Transformed cells were grown in 2XYT media containing 100 μg/ml of ampicillin and 34 μg/ml of chloramphenicol, with protein expression induced by autoinduction(51) during growth at 23°C for 24 hours.

Following expression, cells were lysed by ultrasonication (500 W, 30% duty cycle, 2 seconds on, 10 seconds off for 2 minutes) in cold start buffer (20 mM Tris, 150 mM NaCl, 10 mM imidazole, pH 8) and clarified by centrifugation (28000 x G for 30 minutes at 4°C). Both mseGFP and roGFP2 were purified by NiNTA chromatography (QIAGEN) over a column bed containing 5ml of resin equilibrated with start buffer. The bound proteins were washed three times in wash buffer (20 mM Tris, 150 mM NaCl, 30 mM imidazole, pH 8) followed by elution in elute buffer (20 mM Tris, 150 mM NaCl, 200 mM imidazole pH 8). The N-terminal polyhistidine tag was removed by overnight dialysis (20 mM Tris, 150 mM NaCl, 1 mM beta-mercaptoethanol) with TEV protease at 4°C. The dialyzed protein was again purified by NiNTA chromatography, collecting the flow through. This was then diluted by the addition of two volumes of salt-free buffer (20 mM Tris, 1 mM beta-mercaptoethanol, pH 8), loaded onto a HiTrap Q HP column (Cytiva) equilibrated with 20 mM Tris, pH 8, and eluted using a gradient from 0.05-1M NaCl. Fractions containing proteins were concentrated by ultrafiltration and purified by size exclusion on a Sephacryl S200 column (Cytiva) in SEC buffer (20 mM HEPES, 150 mM NaCl, 1 mM TCEP, pH 7). The proteins typically eluted in two peaks, with fractions from the later peak used for our experiments.

### Determination of mseGFP and roGFP2 fluorescence properties

We measured the absorbance and fluorescence emission spectra of mseGFP and roGFP2 in the same buffer used for SEC (20 mM HEPES, 150 mM NaCl, 1 mM TCEP, pH 7). For absorption, we used a BioSpectrometer (Eppendorf). Fluorescence excitation-emission spectra were collected using a Fluoromax-3 (HORIBA) fluorometer. Absolute concentrations were determined using the absorption of the chromophore in 0.1 M NaOH, which for GFP has an extinction coefficient of 44000 M^-1^cm^-1^ (52). The results of our experiments revealed quantum yield and emission spectra for mseGFP and reduced roGFP2 that were indistinguishable from those of eGFP(53).

### Calculation of FRET radii for transition metals

FRET efficiency depends on the distance between the donor and acceptor molecule and the Förster radius (R_0_) as described by the following formula:

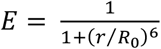

We calculated R_0_ (the distance at which FRET efficiency is 50%) using established methods(31) and assuming an orientation factor *κ*^2^ of 2/3. We estimated the refractive index n as 1.3348, the value described for PBS(54). For determination of spectral overlap with GFP fluorophores, we measured the absorbance of metal complexes using a BioSpectrometer (Eppendorf). When taking these measurements, we attempted to determine the absorbance of metals in complex with thiol-based chelators such as dimercaptosuccinic acid (DMSA) and free cysteine. However, these readings were unstable, and we used EDTA complexes as an approximation instead.

### Fluorescence plate reader assays

For fluorescence assays, 200 μL reactions were set up in individual wells of black 96-well microplates (BRAND) containing 30 nM of fluorescent protein (roGFP2 or mseGFP) and the indicated dilution of each metal or hydrogen peroxide. Titrations were performed in reducing assay buffer (20 mM HEPES, 150 mM NaCl pH 7). We observed that millimolar quantities of reducing agents formed insoluble complexes with some metals; consequently, all protein stocks (which contain TCEP) were diluted >1000 fold for each assay. Experiments involving glutathione included the indicated amount of reduced glutathione, freshly prepared as a 1M stock in water and equilibrated to pH 7 with sodium hydroxide. All reactions were allowed to equilibrate for 30 minutes at room temperature. Readings were performed on a Cytation 5 (BioTek) multimodal plate reader at room temperature. Monochromators were set to either 400 or 485 nm with slit widths of 20 nm for excitation, and 528 nm with a slit width of 20 nm for emission. Samples exhibiting visible precipitation or nonspecific quenching due to light absorbance were excluded from analysis. Binding for samples excepting Fe^2+^ and Fe^3+^ was measured by a ratiometric change in fluorescence indicated by the formula F(λ_x400_, λ_m528_) / F(λ_x485_, λ_m528_). Binding for Fe^2+^ and Fe^3+^ was measured by the total change in F(λ_x488_, λ_m528_) as a result of quenching. Binding for Cu^2+^ was evaluated by ratiometric fluorescence change for roGFP2 and by quenching for mseGFP. Binding data was averaged across experimental days and protein preparations and fitted with a total binding model using Prism 9 (Graphpad).

### Crystallography

Initial crystal screening was performed using the Crystal Screen HT (Hampton) with 20 mg/ml mseGFP in SEC buffer (20 mM HEPES, 150 mM NaCl, 1 mM TCEP, pH 7). Screens were also performed with mseGFP supplemented with 1 mM As^3+^, Pb^2+^, Cd^2+^, phenylarsine oxide (PAO), or 1 mM As^3+^ supplemented with 3 mM glutathione. After optimization, only mseGFP supplemented with PAO produced diffracting crystals. Crystals grown in 0.2 M Li2SO4, 0.1 M Tris pH 8, 32% PEG 3350 were used for cryopreservation and data collection. X-ray diffraction data was collected on a Rigaku R-Axis equipped with HTC plate reader (Rigaku) and processed using HKL2000 (HKL Research). Crystals conformed to the cubic space group I23. Phases were solved by molecular replacement using PHASER-MR(55) with roGFP2 (PDB 1JC0) as the template. Model refinement and the generation of an iterative build OMIT map(41) were performed using Phenix(56) and Coot(57).

### Cell imaging assays

HEK293T cells were cultured in standard culture media (DMEM, 10% FBS, 1X penicillinstreptomycin) (Gibco). Cells grown to >90% confluence in 35 mm plates were transfected with pcDNA3-mseGFP linearized with MluI using Lipofectamine 2000 (Thermo) according to the manufacturer’s instructions. One day post-transfection, cells were removed by treatment with trypsin (0.05%) and plated at 1:10 dilutions in the wells of a 6-well plate containing culture media with dilutions of G418 at concentrations ranging from 0-500 μg/mL. Cells were evaluated for fluorescent colony formation at periodic intervals up to 10 days post transfection. Discrete colonies were selected from wells with G418 concentrations of 300 μg/mL. Colonies were isolated following brief treatment by trypsin and expanded in culture media containing 200 μg/mL G418. For assays involving glutathione depletion, buthionine sulfoximine (Sigma) was added to achieve a final concentration of 100 μM and incubated for 16 hours before data collection.

For imaging experiments, stably transfected cells were incubated in HBSS (Gibco) before adding the indicated amount of metal. Fluorescence imaging was collected over a time using a Nikon A1R inverted microscope with 20X objective and excitation wavelengths of 405 nm (60 power, 125 gain) and 488 nm (11 power, 65 gain) paired with emissions recorded between 520-600 nm and −1 PMT offset. Images were captured using a Galvano scanner, and ratiometric images were collected and analyzed using NIS Elements (Nikon). Ratiometric fluorescence data was exported as LUT histograms for data analysis. Further data analysis was performed in Prism. Histograms for each experimental condition (n≥3) were fitted with a Gaussian function, and the mean values were plotted as the average fluorescence ratio for each time point. We then plotted the resulting timecourse data with a Gompertz growth function.

## Supporting information

Raw and Normalized Data

## Data availability

Research data is included as supporting information in this manuscript. The structural model of mseGFP-PAO is deposited with the Protein Data Bank (PDB 8DTA).

## Supporting information

This article contains supporting information, including raw data and supplementary figures.

## Funding and additional information

This project was funded to J.R by the Competitive Medical Research Fund (CMRF) awarded by the University of Pittsburgh School of Medicine, and to A.C. from startup funds from the University of Pittsburgh. Research in the Carlson lab was also supported by R01GM125638.

## Conflict of interest

The authors declare that they have no conflicts of interest with the contents of this article.

**Table 1.** Data collection and refinement statistics.

|  |  |
| --- | --- |
| Resolution range | 33.89 - 1.811 (1.875 - 1.811) |
| Space group | I 2 3 |
| Unit cell | 126.802 126.802 126.802 90 90 90 |
| Total reflections | 542477 (53313) |
| Unique reflections | 30947 (3068) |
| Multiplicity | 17.5 (17.4) |
| Completeness (%) | 99.71 (100.00) |
| Mean I/sigma(I) | 33.07 (3.83) |
| Wilson B-factor | 23.40 |
| R-merge | 0.1114 (1.018) |
| R-meas | 0.1148 (1.049) |
| R-pim | 0.02724 (0.2504) |
| CC1/2 | 0.999 (0.85) |
| CC* | 1 (0.959) |
| Reflections used in refinement | 30866 (3068) |
| Reflections used for R-free | 1989 (198) |
| R-work | 0.1620 (0.2180) |
| R-free | 0.1813 (0.2684) |
| CC(work) | 0.970 (0.889) |
| CC(free) | 0.965 (0.830) |
| Number of non-hydrogen atoms | 2172 |
| macromolecules | 1868 |
| ligands | 95 |
| solvent | 250 |
| Protein residues | 234 |
| RMS(bonds) | 0.013 |
| RMS(angles) | 1.24 |
| Ramachandran favored (%) | 99.13 |
| Ramachandran allowed (%) | 0.87 |
| Ramachandran outliers (%) | 0.00 |
| Rotamer outliers (%) | 0.49 |
| Clashscore | 1.58 |
| Average B-factor | 28.98 |
| macromolecules | 27.72 |
| ligands | 35.49 |
| solvent | 36.97 |

Statistics for the highest-resolution shell are shown in parentheses.

