## Supplemental Figure 1 for "Studying metal-protein interactions using fluorescent protein indicators"

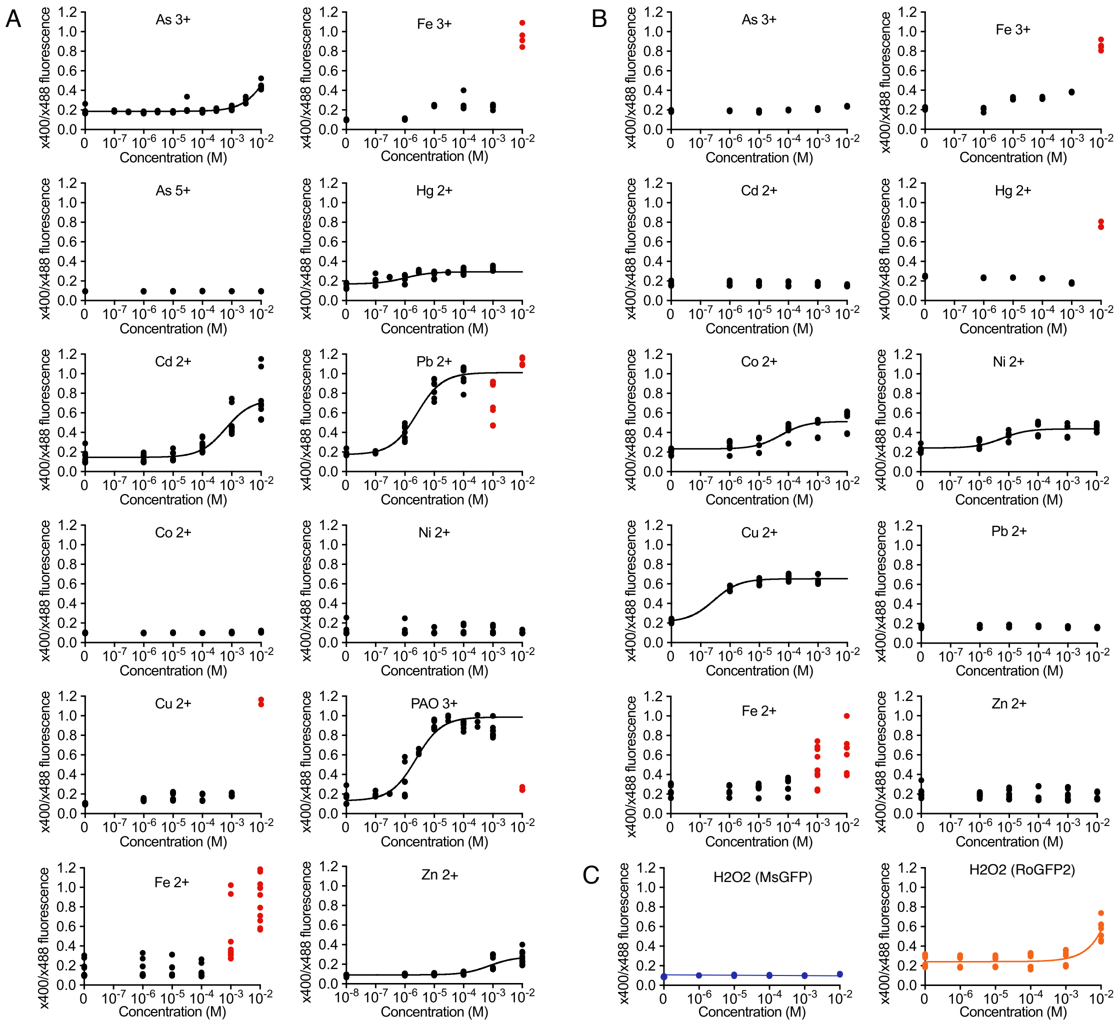


**Supplemental Figure 1. Interactions of msGFP and roGFP2 with metals and oxidizers.** *A*, plot of all titration points with the indicated metals and msGFP. Excluded data points are indicated in red. The remaining data was used to calculate binding curves where shown. *B*, same as for (*A*) but with roGFP2. *C*, titration of $H_2O_2$ produces a ratiometric change on roGFP2 fluorescence but has no effect on msGFP.

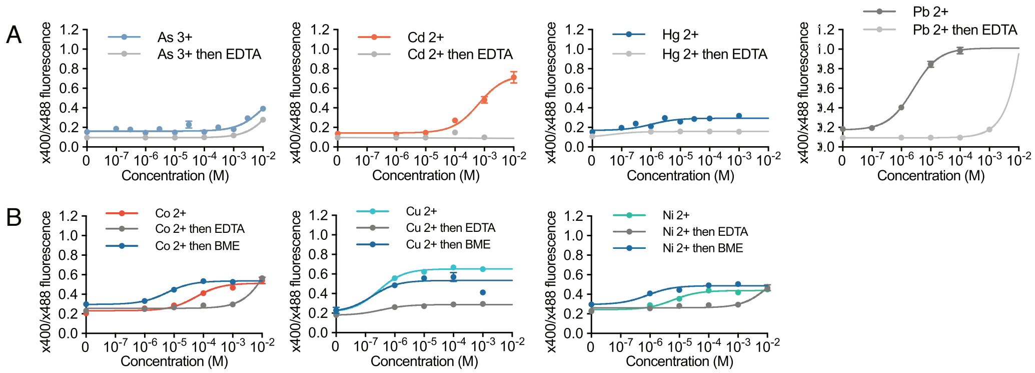


**Supplemental Figure 2. Metal-induced fluorescence changes are reversible by EDTA.** *A*, titrations of metals with msGFP before and after the addition of 5 mM EDTA. *B*, titrations of metals with roGFP2 before and after the addition of 5 mM EDTA or 5 mM β-mercaptoethanol.

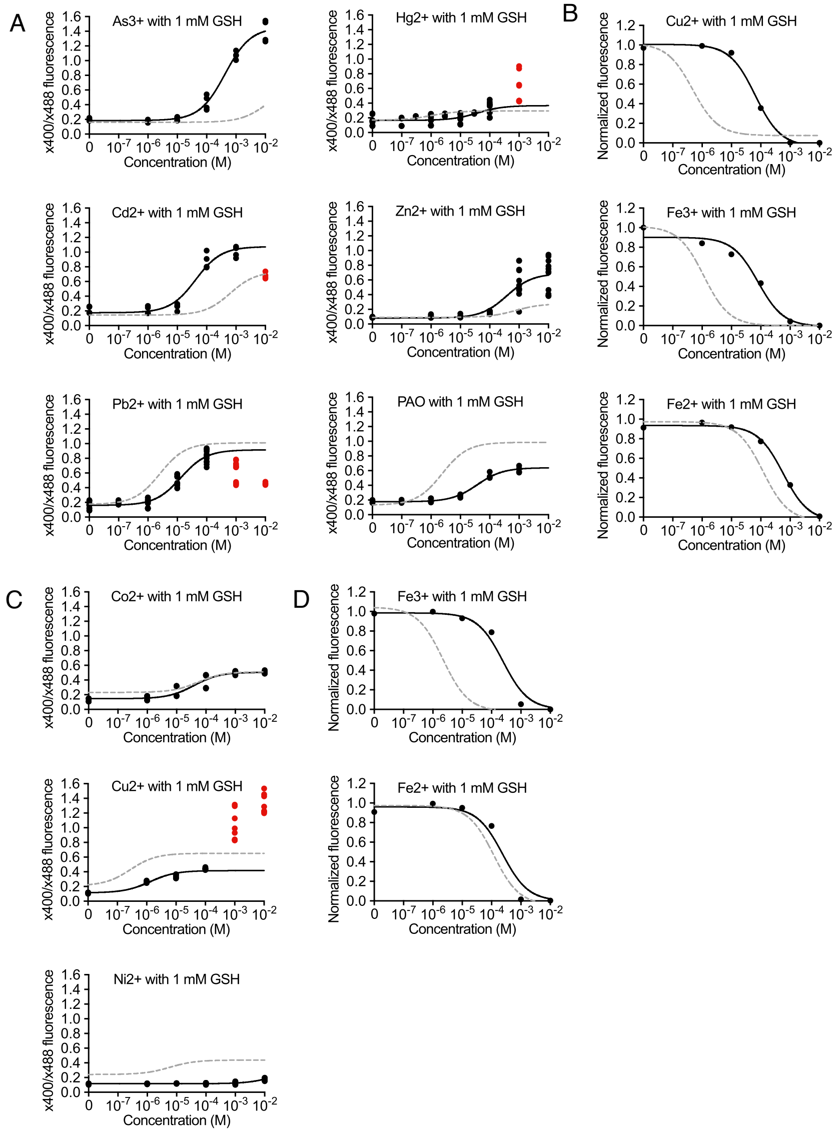


**Supplemental Figure 3. Glutathione effects on metal-reporter interactions.** *A*, ratiometric fluorescence changes in msGFP during titrations of metals in the presence of 1 mM GSH. Excluded data points are indicated in red. Binding curves without glutathione are shown as light grey dashed lines. *B*, quenching of msGFP fluorescence during titrations of metals in the presence of 1 mM GSH. Quenching curves without glutathione are shown as light grey dashed lines. *C*, same as (*A*) except for roGFP2. *D*, same as (*B*) except for roGFP2.
